# CRISPR knockdown of GABA_A_ alpha3 subunits on thalamic reticular neurons enhances deep sleep

**DOI:** 10.1101/2020.12.15.422912

**Authors:** David S. Uygun, Chun Yang, Elena R. Tilli, Fumi Katsuki, James T. McKenna, James M. McNally, Ritchie E. Brown, Radhika Basheer

## Abstract

Identification of mechanisms which increase deep sleep could lead to novel treatments which promote the restorative effects of sleep. Here, knockdown of the α3 GABA_A_-receptor subunit from parvalbumin neurons in the thalamic reticular nucleus using CRISPR-Cas9 gene editing increased the thalamocortical delta oscillations implicated in many health-promoting effects of sleep. Inhibitory synaptic currents were strongly reduced *in vitro*. Effects were most pronounced in the mouse sleep (light) period. Further analysis identified a novel deep-sleep state in mice prior to NREM-REM transitions which was preferentially affected by deletion of α3 subunits. Our results identify a functional role for GABA_A_ receptors on Thalamic Reticular Nucleus neurons and suggest antagonism of α3 subunits as a strategy to enhance deep sleep.

**One Sentence Summary:** Selective genetic knockdown of the major α subunit of GABA_A_ receptors present in the thalamic reticular nucleus enhanced deep sleep in mice.

## Main Text

Sleep is vital for maintaining physical and mental well-being. In particular, thalamocortical delta (0.5-4 Hz) oscillations present in deep non-rapid-eye-movement (NREM) sleep are implicated in a wide range of processes beneficial to health including synaptic homeostasis, cellular energy regulation, clearance of toxic proteins, cognitive performance and mood (*1*–*4*). Conversely, insomnia, traumatic brain injury, obstructive sleep apnea, and other brain disorders are associated with interrupted/fragmented sleep, reduced deep NREM sleep and decreased delta wave power (*5*–*8*). Although hypnotic agents which potentiate the activity of GABA_A_ receptors promote sleep induction, they also reduce delta oscillations, suggesting that a subset of GABA_A_ receptors prevents deep restorative sleep (*9*, *10*). Thus, identification and elimination of this confounding effect of the most widely used hypnotics, which target GABA_A_ receptors, could be beneficial in developing sleep drugs which boost the positive effects of sleep. Recent work showed the thalamic reticular nucleus (TRN) plays a role in promoting delta power (*11*–*13*) and NREM sleep (*14*–*16*). Here we used state-of-the-art CRISPR-Cas9 gene editing to test the hypothesis that GABA_A_ receptors on TRN neurons suppress NREM sleep delta oscillations.

In the adult brain, most GABA_A_Rs consist of two α subunits (α1–6), two (β1–3) subunits (β, and one γ subunit (γ1–3)(*17*) (**Fig. 1a**). The α subunit is a necessary major structure of the GABA_A_R, required for assembling a functional receptor (*18*). In the mouse thalamus, all synaptic GABA_A_Rs in thalamocortical relay nuclei contain α1, whereas GABA_A_Rs in the TRN contain α3 (*19*). To introduce a brain region and cell-type-specific ablation of the α3 subunit gene, we first generated mice which expressed the Cas9 endonuclease in the major subset of TRN neurons which contain the calcium-binding protein parvalbumin (PV) by crossing PV-Cre mice with Rosa26-Lox-stop-lox-Cas9-GFP mice to produce PV-Cas9-GFP offspring. Next, we analyzed the gene sequence of the α3 subunit and selected three loci close to the start codon as target regions expected to maximize CRISPR-Cas9 mediated ablation (**Fig. 1a**). Our target regions were within the extracellular domain (**Fig. 1a**), the major ligand binding component, forming parts of the GABA binding site and the benzodiazepine binding site (*20*). We then constructed an adeno-associated viral (AAV) vector to target the α3 subunit (AAV5-α3-sgRNA-mCherry) by introducing the sequences for the single-guide RNAs (sgRNAs) into an AAV vector plasmid, each driven by the U6 promoter paired with mCherry as a red fluorescent marker (**Fig. 1B**). To test the effect of α3 knockdown (α3KD) on sleep and spectral activity, we recorded cortical oscillations using frontal electroencephalographic (EEG) electrodes and nuchal muscle electromyographic (EMG) electrodes before and after we introduced AAV-α3-sgRNA-mCherry into the TRN via chronically implanted guide cannulae (**Fig. 1B**).

**Figure 1.**
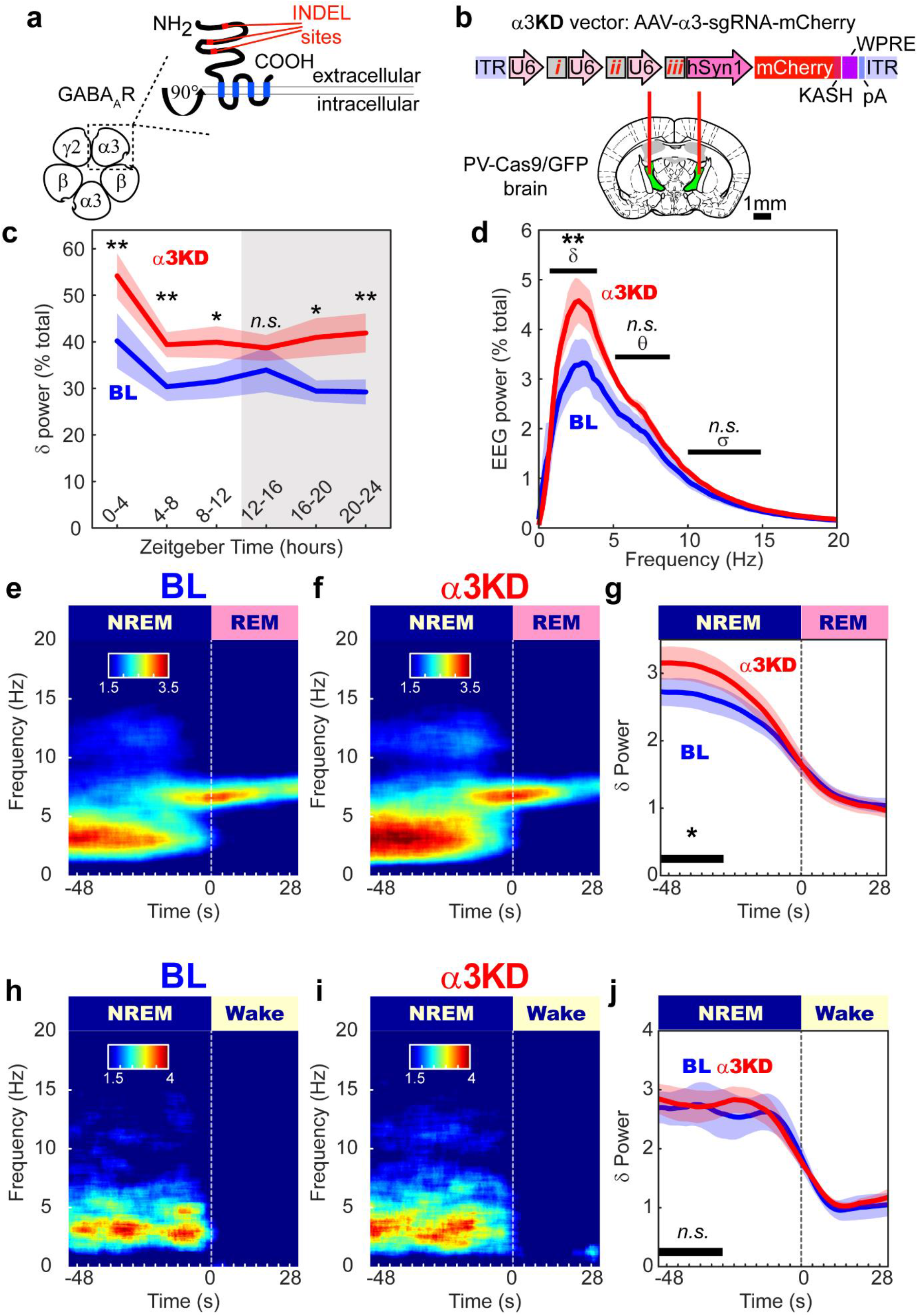
α 3KD in PV+TRN neurons increased NREM delta power, especially at NREM to REM transitions. **a.** The GABA_A_R is a pentameric heteromeric ion channel. CRISPR-Cas9 abscission was directed to three locations (INDEL sites) of the gene which correspond to the large extracellular region of the α3 subunit, a necessary structural as well as ligand binding component in GABA_A_Rs of the TRN. **b.** Adeno-Associated Viral vectors encoding three separate single-guide RNAs (*i, ii & iii*), each driven by its own U6 promoter, and the marker protein mCherry driven by the human synapsin promoter, were injected into the TRN region of PV-Cas9/GFP mice *in vivo* via guide cannulas. **c.** Compared with their baseline (BL) recordings, α3KD mice had higher NREM delta power, with the exception of the first part of the dark period when mice are mostly awake. **d.** Only NREM delta (δ) power is significantly different between BL and α3KD records, theta (θ) and sigma (σ) power were not affected, shown here is the power profile from NREM in ZT-0-4 where we saw the largest effect. **e.** Baseline time-frequency power dynamics presents a surge in delta power in NREM leading to a transition to REM. **f.** After α3KD, the delta power surge in NREM before a transition to REM was increased. **g.** Compared with their BL levels (blue), α3KD mice (red) had higher delta power in the NREM before a transition to REM (p=0.036). **h.** BL time-frequency power dynamics presents a surge in delta power in NREM leading to a transition to wake as well. **i.** α3KD did not increase this delta power surge that occurs during NREM before a transition to wake. **j.** Compared with BL (blue), α3KD (red) did not correspond to a change in delta power in the NREM before a transition to wake (p = 0.47). \*\**p*<0.01, \**p*<0.05, *n.s.* indicates not significant.

α3KD in TRN PV neurons resulted in a marked increase in NREM sleep delta oscillations, which was most pronounced at the beginning of the mouse sleep period (ZT0-4, 47.5 ±21.8 % increase, *t* (5) = −5.159, *p* = 0.0036; **Fig. 1c,** whole 24 hr period shown in **Extended Data Fig. 1**). 6/6 mice with > 85% transduction of TRN PV neurons showed this effect (**Extended Data Fig. 1**). The increase in delta power was similar in magnitude to the increase in delta observed in the first four hours following six hours of sleep deprivation, even though the mice here were not sleep deprived (*21*). We found no change in any other frequency bands in NREM (**Fig. 1d**), and no changes in any frequency bands of wakefulness or REM sleep (**Extended Data Fig. 2**). Only modest changes in the amount of NREM sleep itself were observed, with more NREM only in the first four hours of the dark period when mice are mostly awake. (BL 23.2 ± 2.4 % vs KD 32.4 ± 2.8 %, *t* (5) = −4.7472, *p* = 0.005). We observed a significant reduction in the proportion of shortest bout durations (**Extended Data Fig. 3c**), suggesting more consolidated NREM sleep following α3KD. REM sleep was unchanged. Analysis of sleep spindles using a recently validated algorithm (*22*) did not identify any difference in spindle density, frequency or duration (**Extended Data Fig. 4**). No NREM delta effects were observed in four negative control mice (ZT0-4, *p*=0.19; ZT4-8, *p*=0.38; ZT8-12, *p*=0.5; ZT12-16, *p*=0.39; ZT16-20, *p*=0.95; ZT20-24, *p*=0.67), three of which showed no AAV-α3-sgRNA-mCherry transduction in TRN and one with only 66% transduction of TRN PV neurons.

In humans, delta oscillations are most prominent in the deepest stage of NREM sleep, N3. However, in mice NREM is not generally split into stages. Nevertheless, mouse NREM probably also has degrees of depth which are not evident using standard scoring approaches. In humans, arousal threshold increases with depth of NREM sleep; humans are more likely to awaken from the lighter stages N1 or N2 (*23*). Thus, to separate lighter from deeper NREM sleep in mice, we analyzed delta oscillations at NREM→REM and NREM→wakefulness transitions. The heightened delta power associated with α3KD was only evident during deeper NREM sleep immediately preceding transitions to REM sleep (**Fig. 1d-g**) [means (standard error (SEM): BL = 2.69 (0.22), α3KD = 3.12 (0.24); 17.2 % increase (± 7.7); *t* (5) = 2.27, *p* = 0.036]. No difference was apparent in NREM sleep before transitions to wake (**Fig. 1h-j**). Similarly, no change in delta power was seen during NREM sleep in the initial phase of transitions from wakefulness to NREM sleep (**Extended Data Fig. 5**). We also found the NREM→REM heightened delta was not apparent in the negative control group with either no or poor TRN targeting (**Extended Data Fig. 6**).

Selective deletion of α3 subunits in TRN PV neurons requires the combination of selective expression of Cas9 in PV neurons and sgRNA targeting α3 subunits in the same cells. Furthermore, mCherry (red; marker of sgRNA) was expressed in the majority of TRN PV neurons (green and blue) within the core of the injection site (**Fig. 2a**). In the six α3KD-confirmed mice, we found a high percentage of PV+ TRN neurons (GFP+: 94±1.8%) were transduced by AAV-α3-sgRNA-mCherry, and a large proportion of the TRN (94±1.2%) area was covered (**Fig. 2a**). In preliminary work prior to *in vivo* experiments, we confirmed that Cas9 expression (marked by GFP co-expression) was selective for PV neurons by immunohistochemical staining with PV antibodies (blue secondary antibodies) (**Fig. 2c**), consistent with the previously published validation of Cas9 selective expression in PV+ neurons in this mouse model (*24*).

**Figure 2.**
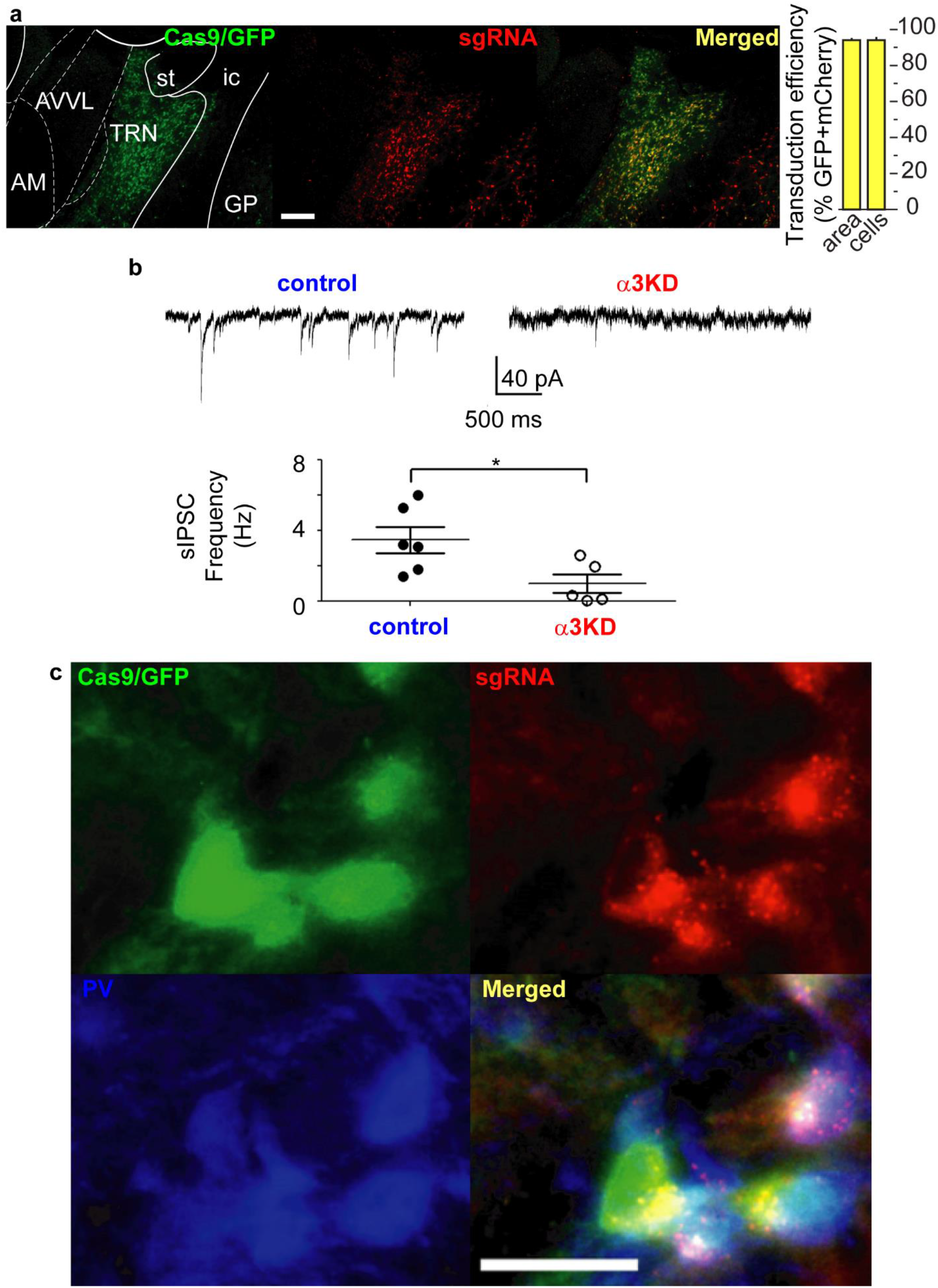
α 3KD in PV+ TRN neurons was validated by histology and *in vitro* electrophysiology. **a.** GFP indicates rich Cas9 expression within the TRN region (green), mCherry reveals widespread transduction of the TRN region by the AAV vector delivering sgRNAs (red) with many of the cells in the area co-expressing both markers (merged; yellow). Scale bar = 200 mm. Percentages of target cells and target area that co-express markers reveal widespread delivery of sgRNAs to target cells in the mice used for *in vivo* studies. **b.** Compared with BL PV+ TRN neurons without KD (left), sIPSCs in α3KD PV+ TRN neurons were significantly reduced (right), with significantly reduced sIPSC frequency (bottom). **c.** High magnification imaging shows triple co-localization of Cas9 (GFP; green), sgRNA (mCherry; red) and PV (immunohistochemical stain; blue), demonstrating successful targeting of PV+ neurons within the TRN. Scale bar = 25 mm.

In a separate group of mice, we verified a functional ablation of GABA_A_ receptors in whole-cell patch-clamp recordings from TRN PV neurons *in vitro*. In control voltage-clamp recordings from TRN PV neurons held at −70 mV in PV-tdTomato mice (which serve as wild type controls with a visual marker of the correct cell phenotype), spontaneous inhibitory postsynaptic currents (sIPSCs) were observed in the presence of glutamate receptor antagonists (20 μM 6-cyano-7-nitroquinoxaline-2,3-dione +50 μM D-(2R)-amino-5-phosphonopentanoic acid). To enhance the driving force for chloride, recordings were made using a patch solution with a high chloride concentration. Thus, IPSCs were detected as inward currents. In PV-Cas9 mice, sIPSCs were significantly reduced in recordings from green (PV-Cas9/GFP) and red (transduced with AAV-α3-sgRNA-mCherry) fluorescent TRN neurons [Frequency: PV-tdTomato: 3.47±0.75Hz (N=6 from four animals); PV-Cas9+AAV-α3-sgRNA: 1.01±0.53 (N=5 from four animals); t(9)=2.560, *p*=0.031, t-test. Amplitude: PV-tdTomato: −38.4±4.6 pA (N=6); PV-Cas9+AAV-α3-sgRNA : −42.1±7.7 pA (N=3 from two animals, in the other recordings sIPSCs were not detectable), t(7)=0.4426, *p*=0.67, t-test] (**Fig. 2b**). Residual sIPSCs in PV+ TRN neurons in PV-Cas9 mice with α3KD retained sensitivity to an α3 selective positive allosteric modulator, TP003 [1μM TP003 increased sIPSC decay time by 13.9±2.1%, t(2)=7.228, *p*=0.02, N=3, paired t-test], suggesting that other α subunits were not upregulated in response to the α3KD.

In another group of mice, we performed the same *in vivo* experimental protocol with a control AAV vector targeting the GABA_A_R α1 subunit. The TRN is devoid of α1 subunits, so this experiment controls for non-specific genetic cutting. In these mice (n=7), we found no change in the amount of NREM delta power (**Fig. 3a),** or other bands in NREM **(Fig. 3b)** following the α1KD. No frequency bands of wakefulness or REM sleep were altered either. The time-frequency analysis at NREM-REM transitions (**Fig. 3c-e**) and at NREM-Wake transitions (**Fig. 3f-h**) showed no changes. Our histologic protocol confirmed a similar degree of targeting success (**Extended Data Fig. S7**).

**Figure 3.**
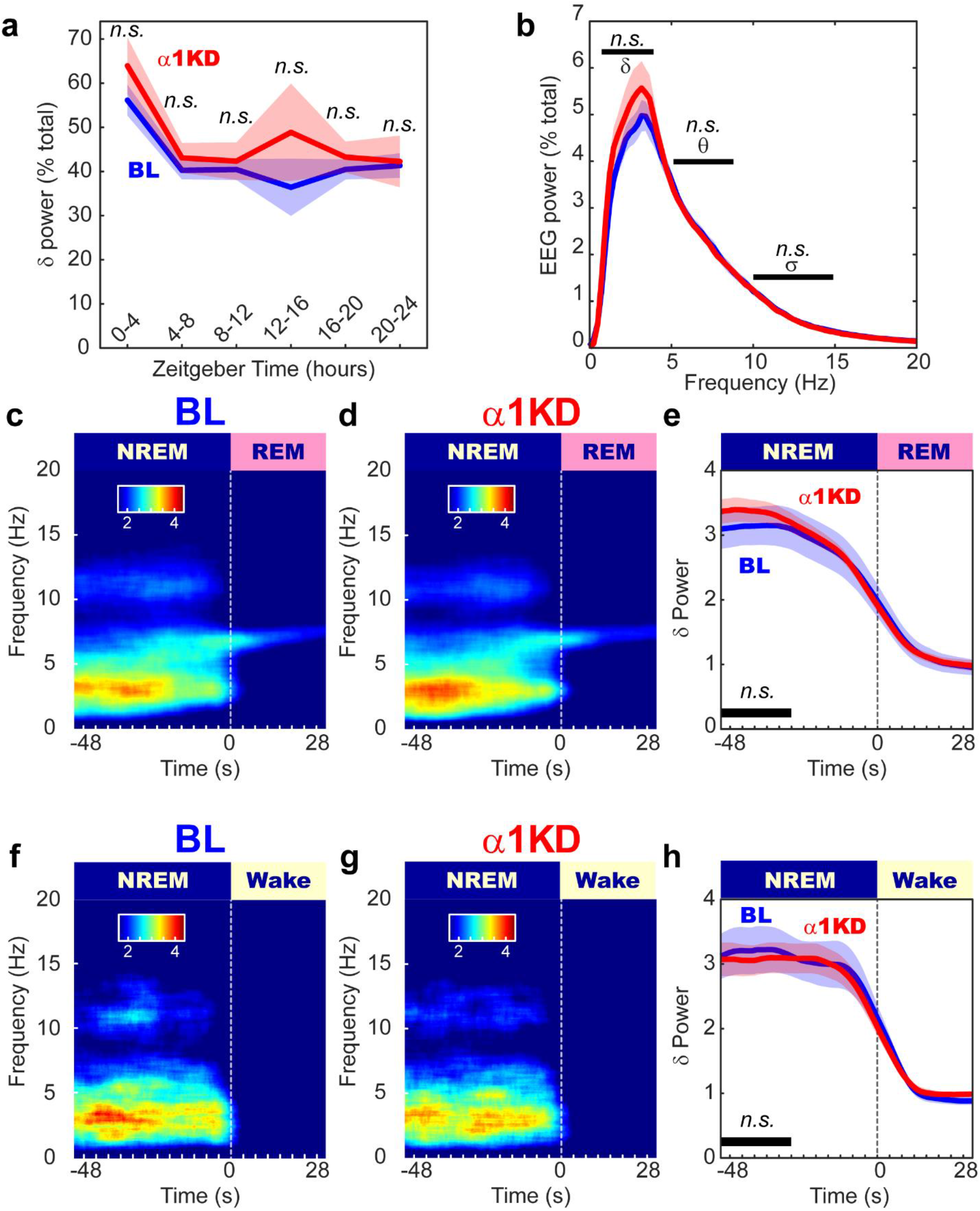
The control cohort with α 1KD in PV+TRN neurons displayed no changes to NREM or wake time, or delta power in any states, including transitions from NREM to REM, in the light inactive period. **a.** Compared with their baseline (BL) recordings, α1KD mice had unchanged NREM delta power. **b.** NREM delta (δ) theta (θ) and sigma (σ) power were not difference between BL and α1KD records, shown here is the power profile from NREM in ZT-0-4 where we saw the largest effect in the α3KD mice. **c.** Baseline time-frequency power dynamics reveals a surge in delta power in NREM leading to a transition to REM. **d.** After α1KD, the delta power surge in NREM before a transition to REM is the same as in baseline records. **e.** Compared with baseline (blue), α1KD (red) mice had unaltered delta power in the NREM before a transition to REM (p = 0.27). **f.** Baseline time-frequency power dynamics reveals a surge in delta power in NREM leading to a transition to wake. **g.** α1KD did not increase the delta power surge in NREM before a transition to wake. **h**. Compared with baseline (blue), α1KD (red) mice had unchanged delta power in the NREM before a transition to wake (p = 0.63). *n.s.* indicates not significant.

In this study, we used cell-type and region-specific CRISPR-Cas9 gene editing *in vivo* to test the functional role of GABAergic inhibition onto TRN neurons in controlling sleep physiology. To the best of our knowledge this is the first time that CRISPR-Cas9 technology has been used *in vivo* to alter thalamocortical oscillations. We found that knockdown of α3-containing GABA receptors, confirmed using *in vitro* recording of sIPSCs, selectively enhances the power of NREM delta oscillations during the sleep period of mice. Our novel analyses identified a deep sleep state prior to NREM-REM transitions which was particularly strongly affected by α3KD. The selectivity of our manipulations was confirmed by our control experiments targeting a closely related subunit, α1, which is not expressed by TRN neurons.

There is an emerging consensus that depolarization of TRN neurons during NREM is an effective way to promote deep sleep (*11*–*16*). TRN neurons receive GABAergic inputs from basal forebrain, lateral hypothalamus and globus pallidus, and several of these neuronal groups maintain a high discharge rate during NREM sleep (*14*, *15*, *25*). Thus, withdrawal of these inputs by removing their postsynaptic targets will lead to a higher discharge rate of TRN neurons during NREM sleep, particularly during deeper stages of NREM prior to REM sleep when excitatory inputs from brainstem aminergic cell groups wane. In turn, increased activity of TRN GABAergic neurons will lead to hyperpolarization of thalamocortical relay neurons, bringing their membrane potential into the correct range to generate delta oscillations. This interpretation of our results is consistent with previous work which suggested that modulating the polarization level and discharge rate of TRN neurons affects delta oscillations and NREM sleep (*11*–*16*).

Our findings differ from previous work which examined constitutive global α3 subunit KO mice (*18*). However, in the constitutive knockout there was no reduction in sIPSC frequency compared to controls in TRN neurons; in fact, there was a modest increase in frequency, plus a significant increase in amplitude, suggesting a developmental compensation. Conversely, we show a significant reduction in sIPSC frequency. Here, the ablation was performed in the adult brain, so developmental compensation was circumvented, which is evident by residual sIPSCs retaining sensitivity to an α3 selective positive allosteric modulator. Therefore, functional ablation of α3 subunits in adults was feasible with the CRISPR-Cas9 approach and, importantly, allowed us to unravel the role of the α3 subunits in sleep-wake patterns and EEG profiles.

In conclusion, CRISPR-Cas9 cell and region-specific gene editing of α3 subunits in adult mice identified a functional role of GABA_A_ receptors on TRN PV neurons in regulating deep NREM sleep. Pharmacological agents which allosterically increase the activity of GABA_A_ receptors containing the α1 or α3 subunits are widely used hypnotics. Unfortunately, they promote light sleep with reduced delta power (*9*, *10*, *26*). Our results suggest that the delta suppressing effect of z-drugs may come from potentiating the α3 containing GABA_A_Rs of the TRN (i.e. the opposite effect that we report here; α3KD leads to increased delta power), a sleep-regulating region which has one of the highest densities of α3 subunits in the brain. Clinically, this is a problem because high NREM delta waves of deep sleep are restorative, important for memory consolidation (*27*) and clearance of toxic metabolites (*3*). Here knockdown of α3 subunits on TRN neurons enhanced deep sleep while not negatively affecting other sleep oscillations or wake power spectra. Mainstay sleep medicines which potentiate GABA_A_ receptors, termed z-drugs, suppress delta waves. Each drug preferentially targets various isoforms. For example, Zolpidem and eszopiclone both bind to α1 α2 and α3, eszopiclone additionally binding α5 (*28*). Interestingly GF-015535-00, not currently in clinical use, is highly selective for α1 and causes far less delta suppression than currently approved z-drugs(*29*). An ideal next-generation hypnotic agent may sedate by positive allosteric modulation of α1 subunits (as do conventional z-drugs); but simultaneously boost delta waves by negatively modulating α3 subunits which are highly expressed in TRN of humans/primates(*30*, *31*), as well as mice. This mechanism could enhance the properties of natural restorative delta oscillations of NREM sleep in manner that current hypnotics do not.

## Author contributions

DSU: Conceptualization, Software, Formal analysis, Investigation, Data Curation, Writing – Original Draft, Writing – Review & Editing, Visualization, Supervision, Funding Acquisition. CY: Conceptualization, Formal analysis, Investigation, Data Curation, Writing – Original Draft, Writing – Review & Editing, Visualization, Funding Acquisition. ERT: Conceptualization, Formal analysis, Investigation, Data Curation, Writing – Review & Editing. FK: Conceptualization, Software, Formal analysis, Investigation, Writing – Review & Editing, Supervision. JTM: Formal analysis, Investigation, Visualization, Writing – Review & Editing. JMM: Conceptualization, Software, Writing – Review & Editing, Supervision, Funding Acquisition. REB: Conceptualization, Resources, Data Curation, Writing – Original Draft, Writing – Review & Editing, Visualization, Supervision, Project administration, FundingAcquisition. RB: Conceptualization, Resources, Data Curation, Writing – Original Draft, Writing – Review & Editing, Visualization, Supervision, Project administration, Funding Acquisition.

## Extended data figure legends

**Extended Data Fig. 1.**
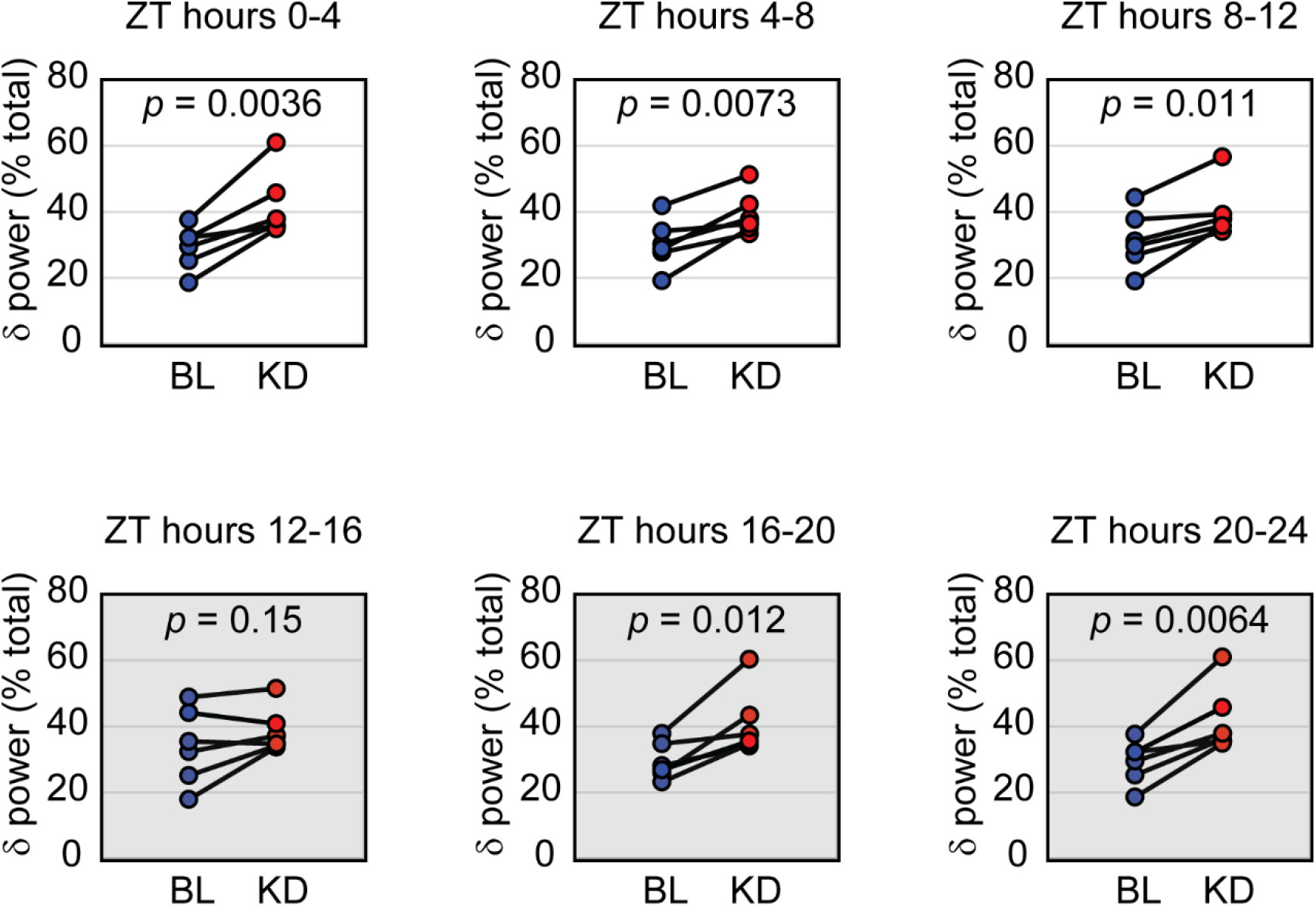
α3KD in PV+TRN neurons consistently increased NREM delta power. Compared with their baseline (BL) recordings, each α3KD mouse had higher NREM delta power in every four-hour period, except the first four hours of the dark period (grey) when mice are mostly awake.

**Extended Data Fig. 2.**
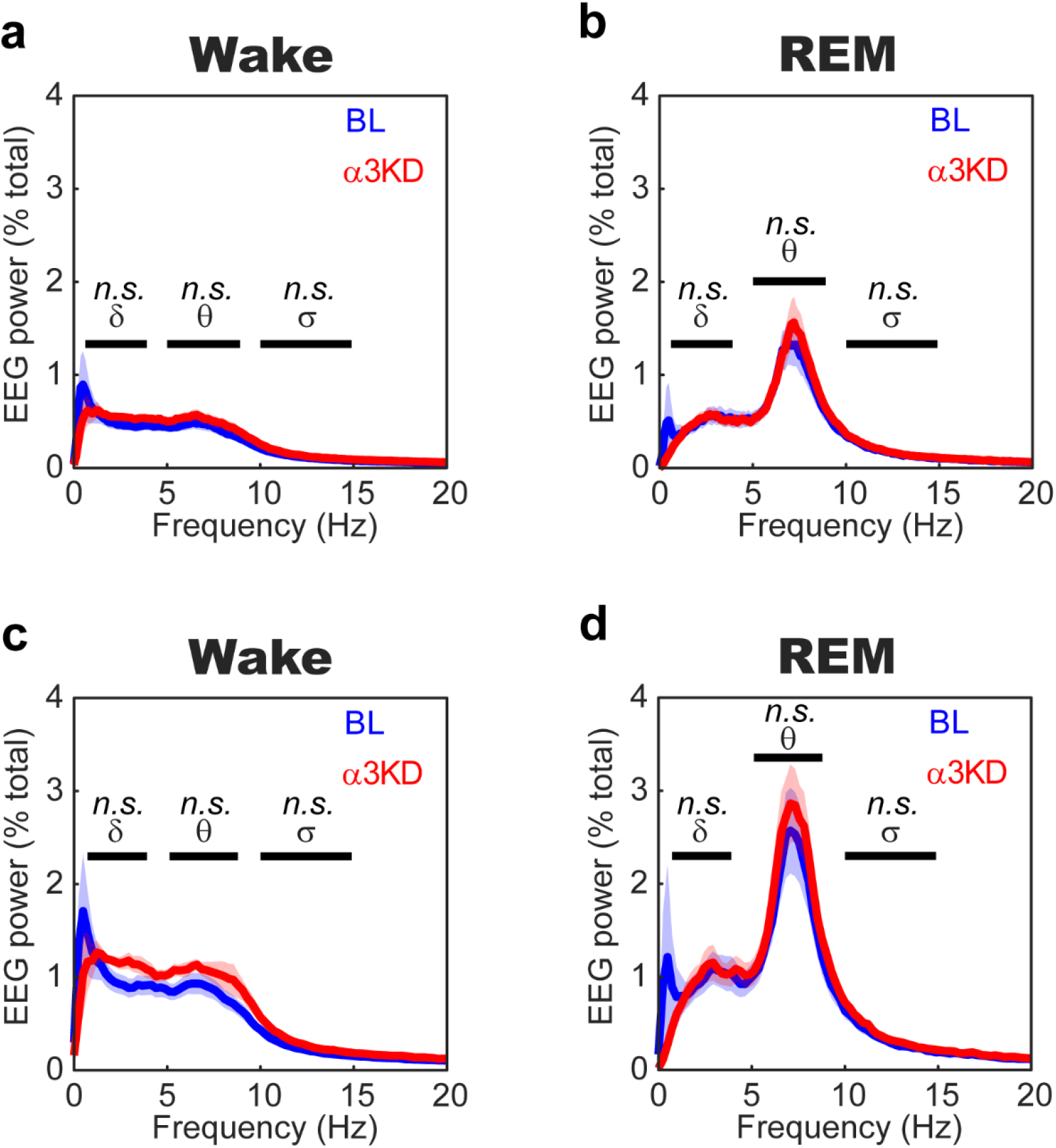
α3KD in PV+TRN neurons did not alter the spectral profile of wake or REM. **a**. Compared with their baseline (BL) levels (blue) from before the α3KD was initiated, α3KD (red) mice (N=6) had the same amounts of delta (δ) power, theta (θ) power and sigma (σ) power in wakefulness during the light period. **b.** Compared with their baseline (BL) levels (blue) from before the α3KD was initiated, α3KD (red) mice had the same amounts of delta (δ) power, theta (θ) power and sigma (σ) power in REM sleep during the light period. **c**. Compared with their baseline (BL) levels (blue) from before the α3KD was initiated, α3KD (red) mice had the same amounts of delta (δ) power, theta (θ) power and sigma (σ) power in wakefulness during the first four hours of the light period. **d.** Compared with their baseline (BL) levels (blue) from before the α3KD was initiated, α3KD (red) mice had the same amounts of delta (δ) power, theta (θ) power and sigma (σ) power in REM sleep during the first four hours of the light period.

**Extended Data Fig. 3.**
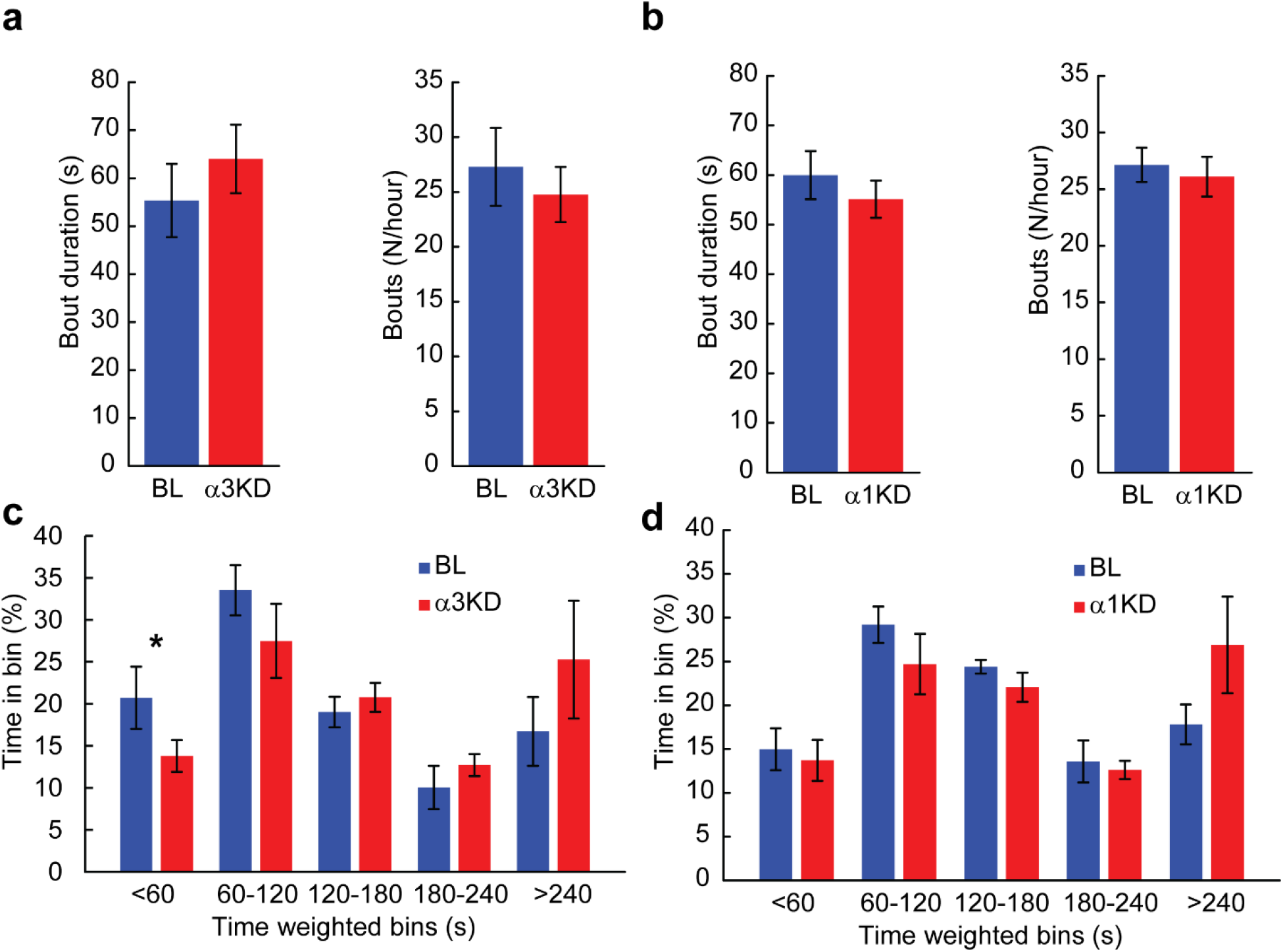
α3KD, but not α1KD, in PV+ TRN neurons decreased time spent in the shortest (<60 s) NREM bouts. **a.** α3KD mice (n=6) did not have altered durations or number of NREM bouts. **b.** α1KD mice (n=7) did not have altered durations or number of NREM bouts. **c.** The proportion of time spent in short bouts lasting <60 seconds was significantly reduced in α3KD mice (p=0.03), but no other time-weighted bins showed any differences between BL and α3KD. **d.** The proportion of time spend in bouts was unchanged in α1KD mice.

**Extended Data Fig. 4.**
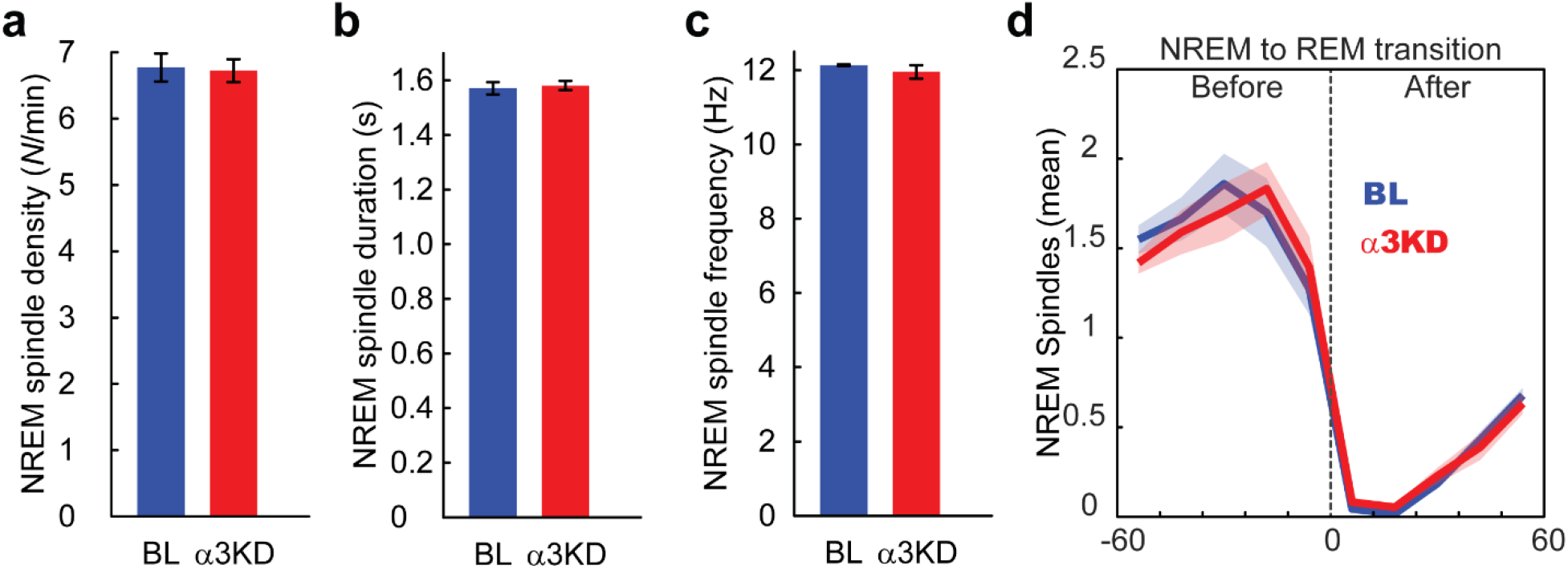
Knockdown of α3 containing GABA_A_Rs in PV+ TRN neurons did not affect sleep spindle activity. **a.** α3KD did not change the density of NREM spindles, as measured by the number of spindles per minute of NREM. **b.** α3KD did not change the duration of NREM spindles. **c.** α3KD did not change the frequency of NREM spindles. **d.** α3KD did not change the number of NREM spindles within the period of NREM before a transition to REM, when sigma power and spindles surge in mice.

**Extended Data Fig. 5.**
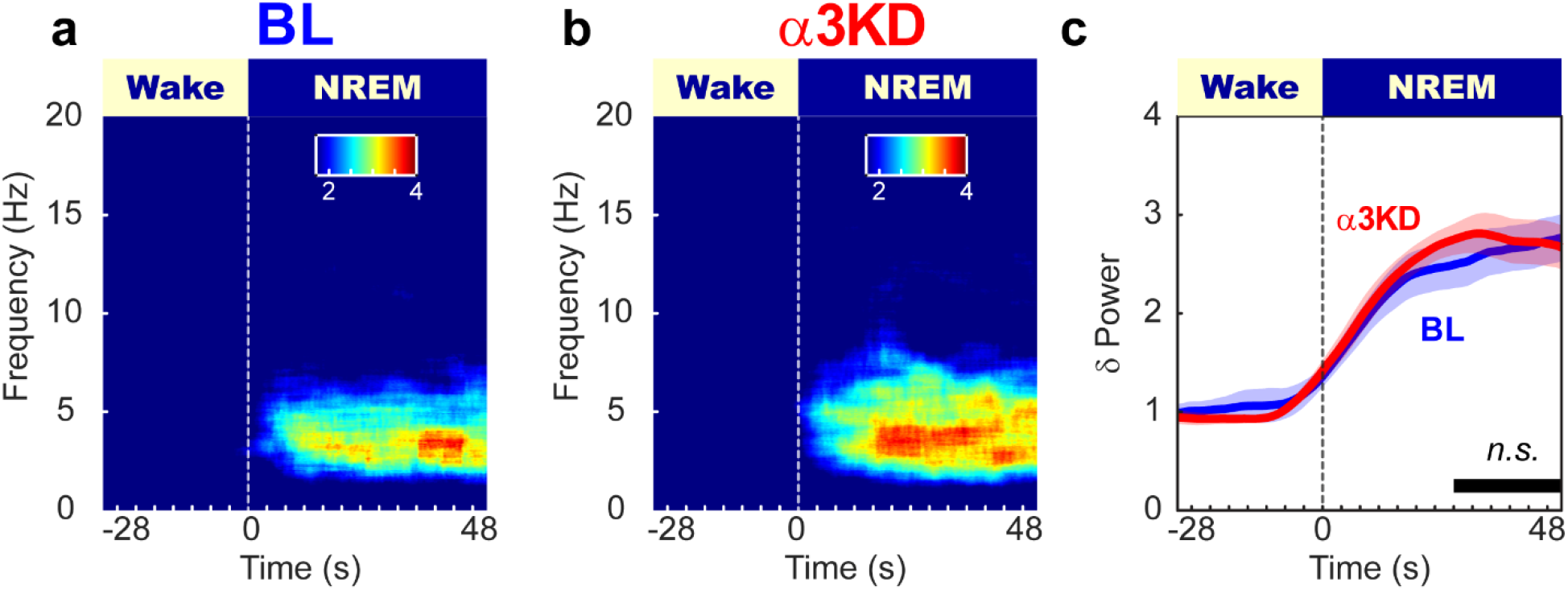
α3KD in PV+TRN neurons did not increase NREM delta power at wake to NREM transitions. **a.** Baseline time-frequency power dynamics presents a surge in delta power in NREM following a transition from wakefulness. **b**. After α3KD, the delta power surge in NREM after a transition from wakefulness appeared similar. **c.** Compared with their BL levels (blue), α3KD (red) mice had similar amounts of delta power in the NREM following a transition from wakefulness (p = 0.28).

**Extended Data Fig. 6.**
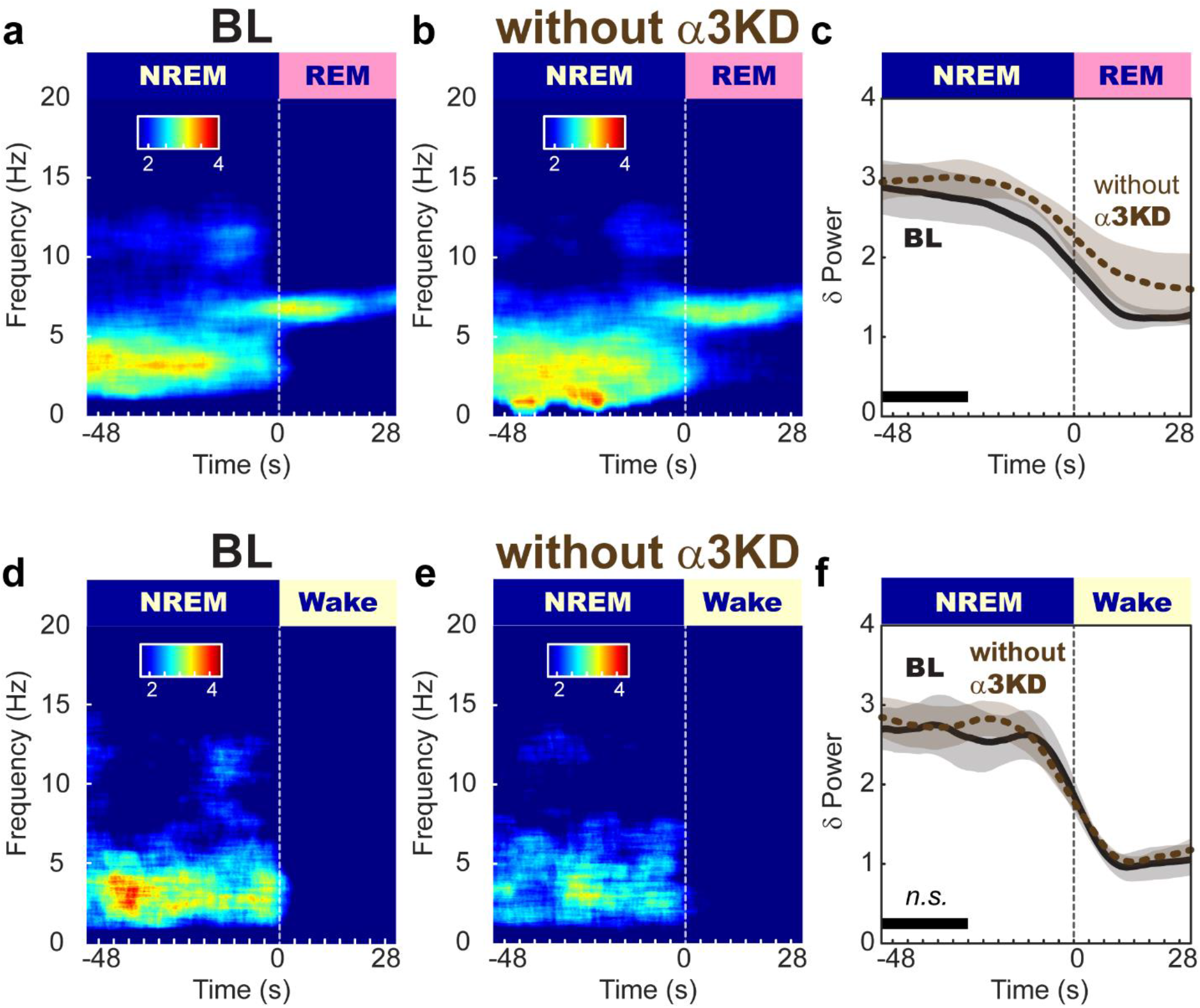
The control cohort with insufficient transduction of PV+TRN neurons, displayed no changes to NREM or wake time, or delta power in any states, including transitions from NREM to REM. **a.** BL time-frequency power dynamics reveals a surge in delta power in NREM leading to a transition to REM. **b.** In the control cohort with no or low transduction of PV+TRN neurons, the delta power surge in NREM before a transition to REM was the same as in their BL records. **c.** Compared with baseline (black), the control cohort with insufficient transduction of PV+TRN neurons (brown) mice had unaltered delta power in the NREM before a transition to REM (p = 0.32). **d.** BL time-frequency power dynamics reveals a surge in delta power in NREM leading to a transition to wake. **e.** the control cohort with insufficient transduction of PV+TRN neurons did not have and increased delta power surge in NREM before a transition to wake. f. Compared with baseline (black), the control cohort which lacked transduction of PV+TRN neurons (brown) mice had unchanged delta power in the NREM preceding a transition to wake (p = 0.9).

**Extended Data Fig. 7.**
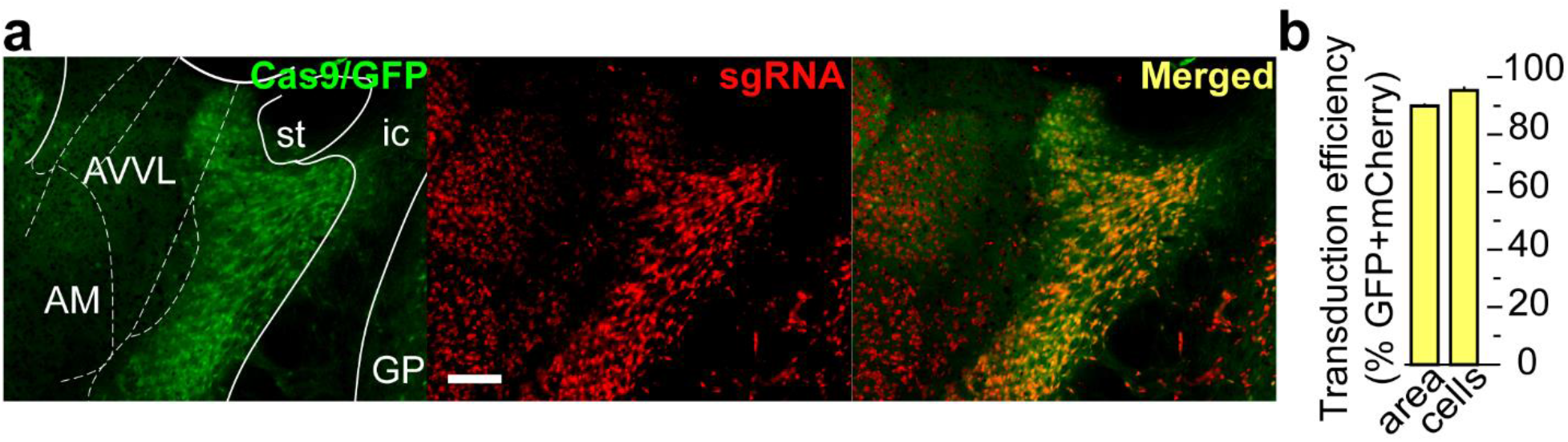
Successful targeting of PV+ TRN neurons by the control vector ‘sgRNA-α1-mCherry’ was validated by histology. **a.** GFP indicates rich Cas9 expression within the TRN region (green), mCherry reveals widespread transduction of the TRN region by the AAV vector delivering α1 targeting sgRNAs (red) with many of the cells in the area co-expressing both markers (merged; yellow). Scale bar = 200 mm. **b.** Percentages of target cells and target area that co-express markers reveal widespread delivery of α1 targeting sgRNAs to target cells in the mice used for *in vivo* control experiments.

## Methods

### Mice

To target our Clustered Regularly Interspersed Short Palindromic Repeats Knock Down (CRISPR KD) selectively to the major subset of Thalamic Reticular Nucleus (TRN) neurons which express Parvalbumin (PV) we crossed male Rosa26-lox-stop-lox-Cas9/GFP (Jackson Labs stock # 026175) mice with female PV-Cre (Jackson Labs stock # 017320) mice, generating mice with the key CRISPR enzyme Cas9 and a green fluorescent protein (GFP) reporter expressed selectively in PV neurons, PV-Cas9/GFP mice. 3-8 month old mice of both sexes were used. No obvious sex differences were observed so data were pooled. For one control group used for *in vitro* sIPSCs recordings, we used PV-tdTomato mice generated by crossing male Rosa26-lox-stop-lox-tdTomato (Jackson Labs stock # 007914) mice with female PV-Cre mice. Mice were housed with a 12h:12h light:dark cycle with lights on at 7am. Food and water were available ad libitum. All experiments were approved by the Institutional Animal Care and Use Committee of VA Boston Healthcare System and conformed to National Institute of Health, Veterans Administration and Harvard Medical School guidelines.

### Adeno-associated viral (AAV) vectors

For this study, the PX552 plasmid described and validated by Swiech et al. (*32*) (Addgene plasmid # 60958; http://n2t.net/addgene:60958; RRID:Addgene_60958) was modified to encode triple U6-sgRNA cassettes each targeting a distinct locus of the target genes. The GFP sequence was replaced by the sequence encoding the red fluorescent protein, mCherry, since GFP was already expressed in PV neurons in the PV-Cas9/GFP mice used for experiments. This newly constructed custom plasmid became our backbone for subsequent design of our control vector.

#### AAV-<u>α3</u>-sgRNA-mCherry

To selectively inactivate the gene encoding the only a subunit (α3) of GABA_A_ receptors expressed in TRN (*19*), we used the above custom-designed AAV vector to deliver three sgRNAs targeting α3 subunit to TRN. α3 GABA_A_ subunit gene (GABRA3; NCBI Reference Sequence: NM_008067.4): *i:* 5’ TCTTCACTAGAATCTTGGAT 3’ *ii:* 5’ GGACCCTCCTCTATACAATG 3’ and *iii* 5’ TTGTTGGGACAGAGATAATC 3’. We found *in silico* that the α3-sgRNAs had no off-targets in the mouse genome (*33*). The CRISPR design tools http://crispr.mit.edu/ and http://chopchop.cbu.uib.no/ (*34*) were used to aid initial identification of possible sgRNA sequences corresponding to the N-terminal domain (**Fig. 1A**). The sgRNA candidate sequences were manually selected by alignment analysis (Bioedit; open source software, Ibis Therapeutics, CA) to confirm that they did not target the closely related α1 subunit gene; (NCBI Reference Sequence: NM_010250.5). Though not found in TRN, α1 is present in the thalamus (*19*) and we wanted to avoid off target GABA_A_ receptor KD, which we viewed as the only tangible confound beyond non-specific CRISPR-Cas9 abscission. All three α3-sgRNA showed minimal homology with α1 and was not indicated to be an off-target site *in silico*.

#### AAV-<u>α1</u>-sgRNA-mCherry

We designed another vector bearing three sgRNAs targeting the α1 gene of GABA_A_ receptor. One sgRNA with low homology to α3 but perfect homology to α1 with a PAM sequence and the other two mock-sgRNAs with perfect homology to non-PAM bearing areas of α1 gene were chosen. *i:* 5’ CTCATTCTGAGCACACTGTC 3’ *ii,* 5’ TTTTTCCGTCAAAGTTGGAA 3’ *iii* 5’ TTGGACAAACAGTTGACTCT 3’. All three sequences from mouse α1 gene showed no off-targets *in silico* in the mouse genome (*33*). We used snapgene software for plasmid design, construction was outsourced to GenScript (New Jersey, United States; www.genscript.com/) and validated by restriction mapping by gel electrophoresis and sequencing. Plasmids, thus validated, were packaged into AAV5 by the University of North Carolina vector core facility. Vector titers (~1-4×10^12^ particles/ml) were determined by dot-blot analysis and used for microinjections into the TRN. (**Fig. 1B**).

### Stereotaxic surgery and AAV microinjections into TRN

Executing the selective CRISPR KD of α3 subunits in TRN PV neurons requires expression of the sgRNAs targeting α3 subunits combined with the Cas9 expressed in PV neurons of the transgenic mice. We achieved this combination by stereotaxic injection of the AAVs expressing sgRNAs into the TRN of the PV-Cas9/GFP mice. Using a Kopf stereotaxic frame, we chronically implanted cannulas bilaterally above the anterior TRN (Plastics One; Connecticut, United States), Part # C315G/SPC, total length 12mm), with retainers inserted (Plastics One, Part # C15I/SPC), at AP −0.7mm, ML ±1.4, DV −1.5. The coordinates for cannula location were selected to be 2 mm above the TRN to allow subsequent microinjection into TRN without extensive damage to TRN. To record cortical electrical activity, bilateral frontal neocortical EEG screw electrodes (Pinnacle Technology Inc.; Kansas, United States; Part # 8403) were placed at AP +1.9 mm, ML ±1.5 with a reference electrode at AP −3 mm, ML +2.7 and a ground electrode at AP −6 mm ML = 0 respectively and soldered to a headmount (Pinnacle Technolology Inc.; Part # 8201-SS). EMG electrodes were placed in the nuchal muscle. All the chronically implanted components were secured with dental cement (Keystone industries, Bosworth Fastray; Part # 0921378).

Following one full week of recovery from cannula/electrode implantation, and collection of baseline EEG/EMG records (see next section), AAV microinjections were made into the TRN. Mice were anesthetized by isoflurane (1.5-4% in O2) and depth of anesthesia was monitored by breathing rate, pedal withdrawal and tail pinch reflexes. A 5 μl Hamilton syringe (Part # 87908, Model 75 SN SYR with 33g cemented needle) loaded with viral solutions was lowered through the cannula, 2 mm beyond the cannula tip, into the TRN (DV −3.5). 1 μl microinjections were delivered at 0.05 μl/minute using a micropump (KD Scientific Legato 130, Massachusetts, United States). Doses of Meloxicam (5mg/kg; intraperitoneal) were given immediately after surgery and again 22-24 hours later, to mitigate any pain associated with the surgery. One month following AAV injection, EEG/EMG signals were again recorded and compared to baseline recordings.

### Electroencephalogram (EEG)/Electromyogram (EMG) recordings

To study sleep wake-states and thalamocortical oscillations, we recorded EEG and EMG using Pinnacle Technology Inc. 3 channel (2 EEG/1EMG) systems for mice (Part # 8200-K1-SL), using its acquisition software (Sirenia Acquisition). Mice were tethered to the system via mouse pre-amplifiers (Pinnacle Technology Inc. Part # 8202-SL) and 24-hour recordings were collected between zeitgeber time 0-24 following a 48-hour period of habituation to the recording apparatus. EEG/EMG data was sampled at 2 kHz, amplified 100x and low pass filtered at 600Hz. In one mouse from the α3KD cohort and one mouse from the control cohort which lacked transduction of PV+TRN neurons, we missed data from ZT0-ZT2. In one mouse from the control cohort which lacked transduction of PV+TRN neurons we missed data from ZT0-ZT3. These were due to software malfunctions. We dealt with this by continuing the recordings into the subsequent day and using the same ZT hours to replace the missing data.

### Sleep-wake scoring and EEG analysis

We manually scored sleep-wake states from EEG and EMG records using four second epochs as follows: Wake was scored when EEG showed a desynchronized low amplitude signal with muscle tone evident by a large EMG signal; the large EMG signal did not need to be phasic in appearance to be characterized as wake. NREM sleep was scored when the EEG signal showed large amplitude, slow synchronized waves, and a low EMG signal, except for very brief bursts which were considered twitching. REM sleep was scored when the EEG signal presented a repetitive stereotyped ‘sawtooth’ signal in the theta range (5-9 Hz), with a nearly flat EMG signal. Artifacts were dealt with as follows: periods that appeared to be wakefulness, but the EEG was contaminated by crosstalk with EMG signals, were scored as ‘wake-exclude’. Periods of NREM or REM with large amplitude DC shifts were extremely rare and labelled as ‘NREM-exclude’ or ‘REM-exclude’. We used all epochs, even those marked ‘exclude’, in behavioral analyses such as time in state or bout analysis, but we only included artifact free data for analyses of EEG signals, such as power spectral density or time-frequency analyses. Scoring was performed in Sirenia Sleep, and EEG signals and scored epochs were exported for further analysis in Matlab.

Power spectral density of wake, NREM and REM was computed using the Matlab pwelch function using an 8-second Hanning window with 50% overlap. To produce time-frequency spectrograms for state-transition analysis, we first down-sampled the data to 40Hz and screened for outliers, replacing values more than ten standard deviations away from the mean with zeros. We then used the multi-taper method (*35*) (Chronux Toolbox; Chronux.org) function (5 tapers with 10 second sliding window in 100 ms steps). Spectra were computed for each state-transition per mouse and mean averaged. Each within-animal mean averaged time-frequency spectrum was then normalized to its average power from wakefulness (I.E. Frequency bin from spectra/Frequency bin from wake). These normalized spectra were then mean averaged across all the animals in the group for a grand mean averaged spectrogram. Mean averaged power in the delta band (0.5-4 Hz) was plotted with standard error envelopes, smoothed using a 5 second moving average window. We analyzed sleep spindles using our validated automated spindle detection algorithm (*22*).

### Histology

Mice were transcardially perfused with 10 ml phosphate buffered saline (PBS) followed by 10 ml of 10% formalin for fixation. Brains were extracted and subsequently immersion-fixed in 10% formalin for 1-2 days, followed by30% sucrose solution in PBS before tissue was sliced at a thickness of 40□m on a freezing microtome (Leica Biosystems, Illinois, United States).

We first confirmed previous findings (*24*) that Cas9 was expressed selectively in TRN PV neurons, by performing immunohistochemistry for PV in coronal sections containing TRN from PV-Cas9/GFP mice. Free-floating sections in wells were treated with PV primary antibodies (rabbit anti-PV; 1:200; AB11427; Abcam, Cambridge, MA) and blue secondary antibodies (donkey anti-rabbit IgG conjugated to AlexaFluor 405; 1:100; ThermoFisher Scientific; Z25313).

Sections were mounted on microscope slides and coverslipped using Vectrashield Hard Set mounting medium (Part # H-1400, Vector Labor). Images were collected on a Zeiss Image2 microscope, outfitted with a Hamamatsu Orca R2 camera (C10600) and Stereo Investigator software (MBF Bioscience). For our *in vivo* sleep-wake experiments, we visually confirmed the presence and quantified co-expression of sgRNAs and Cas9 within TRN by identifying fluorescent markers of sgRNAs (mCherry) and Cas9 (GFP). GFP+ neurons were identified by green fluorescence in the cytoplasm (excitation:emission 488:509) and mCherry+ neurons were identified by red fluorescence in the cytoplasm (excitation:emission 590:617). We also performed immunohistochemistry for PV, as described above, in tissue from PV/Cas9GFP animals injected with AAV sgRNA, to demonstrate sgRNA transduction (mCherry) specific to Cas9+ neurons (GFP) that are PV+ (blue AlexaFluor 405) in TRN (Figure 2E).

For measures of transduction success, we calculated percentages of targets based on an average of two sections per brain, except for one brain where only one section showed transduction. mCherry signals marking AAV transduction were consistently found within the anterior-posterior location, Bregma −0.8±0.2 mm. Target areas were measured using the Fiji free-hand tool (https://imagej.net/Fiji) which reports a manually drawn area in pixels. We first drew an area around the GFP+ TRN region; within this, we drew a second area which was mCherry+. Percentages reported represent the percent of the mCherry+ area divided by the total GFP+ area. Targeted cells were counted manually using the Fiji multipoint tool. We first counted all the GFP+ cells within the TRN region, then we counted all the cells that were both mCherry+ and GFP+ within that region. Percentages represent the number of double labelled (mCherry+GFP+) cells divided by total number of GFP+ cells within the TRN. Mice were considered successful cases if any TRN region was found to be positive for both mCherry and GFP. All successful cases were used to quantify transduction, as described above. In three cases, no signal for mCherry was found within the GFP+ TRN region and were excluded from the cohort. The *in vivo* data from these three mice were used only in the serendipitous control cohort of Fig S4.

### *In vitro* slice electrophysiology

Mice were deeply anesthetized with isoflurane and decapitated. Coronal brain sections containing TRN (Bregma −0.46 to −0.94 mm) were cut at 300 μm-thickness with a Leica VT1200S vibratome (Leica Biosystems Inc., Buffalo Grove, IL, USA) at 4°C. After slicing, the slices were placed into ACSF containing the following (in mM): 124 NaCl, 1.8 KCl, 25.6 NaHCO_3_, 1.2 KH_2_PO_4_, 2 CaCl_2_, 1.3 MgSO_4_, and 10 glucose (300 mOsm), saturated with 95% O_2_/5% CO_2_ for at least 1h at room temperature before being transferred to the recording chamber and superfused with warm ACSF (32 °C) at 2-3 ml/min.

Spontaneous inhibitory post-synaptic current (sIPSC) recordings focused on the anterior-dorsal section of the TRN and target cells were identified by GFP expression and mCherry expression. We filled 3-6 MΩ patch pipettes with intracellular solution of the following composition (in mM): 130 KCl, 5 NaCl, 2 MgCl_2_, 10 HEPES, 0.1 EGTA, 2 NA_2_ATP, 0.5 NaGTP, 4 MgATP and 1 spermine, pH 7.25 with KOH (280 mOsm). sIPSCs were recorded at −70 mV in the presence of the glutamate receptor antagonists (20 μM 6-cyano-7-nitroquinoxaline-2,3-dione +50 μM D-(2R)-amino-5-phosphonopentanoic acid) using a Multiclamp 700B amplifier and pClamp 10.0 software (Molecular devices; California, United States). A 1 min period after 5 min application of the glutamate receptor antagonists was used for statistical analysis (Igor software, WaveMetrics, Inc., Portland, OR, USA). Only well resolved events with amplitudes >10 pA were analyzed. Series resistance was 6-20 MΩ and was not compensated. Sampling rate was 20 kHz. Records were low-pass (Bessel) filtered at 1KHz.

### Statistical information

We used two-tailed paired t-tests to compare BL areas vs KD areas under delta power or other defined frequency bands (represented by bold black lines and symbols d, q or s; ‘significant’ or ‘not significant’ is indicated by * or *n.s.* respectively) in NREM, wake or REM. p-values less than Holm-Bonferroni corrected alpha levels were considered significant. We found a unidirectional effect on NREM delta, so we used a one-tailed test in subsequent delta power analyses, comparing areas under dynamic changes in delta power at state-transitions (also marked by a bold black line with ‘significant’ or ‘not significant’ indicated by * or *n.s.* respectively). We used a two-tailed paired t-test in bout analyses and spindle analyses, and an unpaired t-test to compare control vs KD sIPSCs. All statistics were performed in Matlab or Microsoft Excel.

